# In-cell quantitative structural imaging of phytoplankton using 3D electron microscopy

**DOI:** 10.1101/2020.05.19.104166

**Authors:** Clarisse Uwizeye, Johan Decelle, Pierre-Henri Jouneau, Benoit Gallet, Jean-Baptiste Keck, Christine Moriscot, Fabien Chevalier, Nicole L. Schieber, Rachel Templin, Gilles Curien, Yannick Schwab, Guy Schoehn, Samuel C. Zeeman, Denis Falconet, Giovanni Finazzi

## Abstract

Phytoplankton is a minor fraction of the global biomass playing a major role in primary production and climate. Despite improved understanding of phytoplankton diversity and genomics, we lack nanoscale subcellular imaging approaches to understand their physiology and cell biology. Here, we present a complete Focused Ion Beam - Scanning Electron Microscopy (FIB-SEM) workflow (from sample preparation to image processing) to generate nanometric 3D phytoplankton models. Tomograms of entire cells, representatives of six ecologically-successful phytoplankton unicellular eukaryotes, were used for quantitative morphometric analysis. Besides lineage-specific cellular architectures, we observed common features related to cellular energy management: *i)* conserved cell-volume fractions occupied by the different organelles; *ii)* consistent plastid-mitochondria interactions, *iii)* constant volumetric ratios in these energy-producing organelles. We revealed detailed subcellular features related to chromatin organization and to biomineralization. Overall, this approach opens new perspectives to study phytoplankton acclimation responses to abiotic and biotic factors at a relevant biological scale.

## Introduction

Phytoplankton plays a critical role in supporting life on Earth. By converting CO_2_, sunlight and nutrients into biomass and oxygen, these unicellular phototrophs are responsible for about 50% of primary productivity (Field et al., 1998) and contribute significantly to food webs and to the biological CO_2_ pump in the oceans. They are ubiquitous in marine and freshwater ecosystems and include prokaryotes and eukaryotes, the latter having acquired photosynthesis capacity up to 1.5 billion years ago through endosymbiotic events. Eukaryotic phytoplankton encompasses a great diversity of lineages (e.g. diatoms, dinoflagellates, haptophyceae, chlorophyceae, rhodophceae) with different morphologies and sizes (from 0.8 to a few tens of microns) (Not et al., 2012).

So far, phytoplankton morphological features have been mainly visualized by light microscopy and 2D electron microscopy studies (Andersen et al., 2015; Embleton et al., 2003; Rodenacker et al., 2006; Schulze et al., 2013; Sosik and Olson, 2007), often associated with assessment of photosynthetic activity (Hense et al., 2008; Schulze et al., 2013). A high-throughput confocal fluorescence 3D imaging has recently been developed to scan, classify and quantify phytoplankton cells collected in different oceanic regions (Colin et al., 2017). However, optical microscopy studies have insufficient resolution to reveal microstructural features, emphasising the need to develop complementary imaging approaches to study the cellular and subcellular bases of phytoplankton physiology and cell biology. Recently, a few studies (Decelle et al., 2019; Engel et al., 2015; Flori et al., 2017) have highlighted the potential of 3D electron microscopy (EM) imaging to explore these fundamental aspects of phytoplankton. This is a critical aspect, since previous work has suggested, for instance, that the physiology and metabolism of diatoms are determined by their peculiar cell organisations and more particularly by the morphology and arrangement of key energy-producing machineries, such as the plastids and mitochondria (Bailleul et al., 2015; Flori et al., 2017).

Here, we present a complete imaging workflow to access the cellular and subcellular features of phytoplankton at the nanometric scale based on Focused Ion Beam - Scanning Electron Microscopy (FIB-SEM) (Hawes and Hummel, 2015; Narayan and Subramaniam, 2015; Titze and Genoud, 2016). This technique has already been applied successfully to provide 3D models of eukaryotic cells (Decelle et al., 2019; Flori et al., 2017; Gavelis et al., 2019). Although the spatial resolution of FIB-SEM (4-8 nm) is lower than the resolution of transmission electron microscopy (Engel et al., 2015; Wietrzynski et al., 2020), it has the advantage of providing contextual 3D images of whole cells at a specific time. The FIB-SEM technique does not require *a priori* knowledge on proteins and genomes to observe cellular structures, making it applicable where other label-dependent methods (e.g. super-resolution microscopy) are not. We applied FIB-SEM imaging to six monoclonal cultures of different eukaryotic microalgae representing major phytoplankton lineages. We optimised every stage of the workflow (sample preparation, 3D imaging, filtering, segmentation, and 3D modelling) to generate accurate 3D reconstructions of whole cells in their close-to-native states, suitable for ultrastructural inspection and morphometric analysis (surfaces and volumes). These models allowed us to compare and quantify their subcellular features, leading to the identification of conserved structural characteristics. We also observed species-specific subcellular structures related to the organisation of the genome and related to biomineralisation. Overall, these results highlight the value of whole-cell 3D FIB-SEM tomography to study the cellular architecture of phytoplankton and their responses to various abiotic and biotic factors. This approach may be relevant in the context of climate change scenarios (Intergovernmental Panel on Climate, 2014), where modifications of the acclimation capability of phytoplankton could alter the entire trophic chain and biogeochemical cycles in the ocean.

## Results and Discussion

### 1 Optimization of sample preparation and image analysis workflow

We collected microalgal cells maintained in culture from different eukaryotic lineages: Mammiellophyceae (*Micromonas commoda RCC 827*), Prymnesiophyceae (*Emiliania huxleyi RCC 909*), Bacillariophyceae (*Phaeodactylum tricornutum Pt1 8.6*), Pelagophyceae (*Pelagomonas calceolata RCC 100*), Dinophyceae (*Symbiodinium pilosum RCC 4014* clade A), Cyanidiophyceae (*Galdieria sulphuraria*) (Supplementary Table 1).

To minimize artefacts related to sample preparation, living cells were cryofixed with high-pressure freezing followed by slow freeze-substitution and resin embedding (see Materials and Methods). This method provides enough contrast to distinguish and classify subcellular features with electron microscopy and preserves cellular features better than chemical fixation at room temperature. Embedded cells were imaged with FIB-SEM (Figure 1A) in automatic mode with a voxel size of 8 nm, to obtain complete datasets (i.e. stacks of 2D images, Supplementary Videos 1 to 6) with a workable (~1000-1200) number of frames. FIB-SEM datasets were further processed to obtain 3D models using open-access software (Figure 1). Frames were aligned using the template matching plugin implemented in Fiji (Figure 1A). They were filtered to reduce noise while preserving cellular details. Based on the effectiveness in highlighting organelle boundaries, different filters were used for the different microalgae (Figure 1B). A linear (Gaussian) filter followed by edge enhancement (sharpening) was chosen for *Micromonas*, *Emiliania*, *Phaeodactylum* and *Pelagomonas*, while a non-linear filter (median filter) was preferred in the case of *Galdieria* and *Symbiodinium*. These choices reflect the different cellular features and biochemical composition of each taxon (e.g. the presence of a thick cell wall in *Galdieria*), which results in variable contrast.

**Figure 1:**
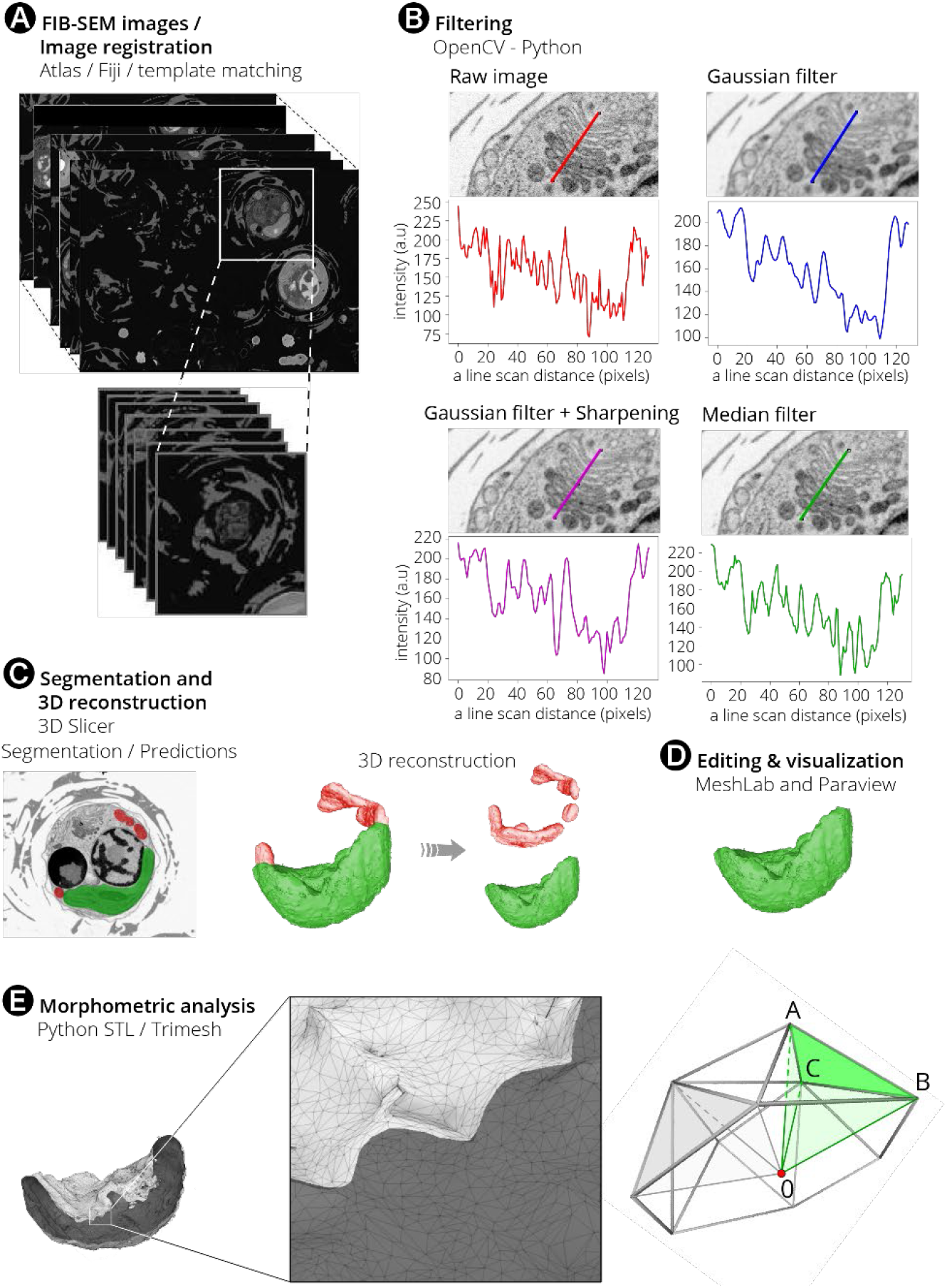
Flowchart of image processing from data acquisition to 3D reconstruction and morphometric analysis. The pipeline includes data acquisition with FIB-SEM and registration with Fiji **(A)**. Single cells are selected from the whole FIB-SEM stack, images are registered before inverting their contrast. Stack are filtered in Python using the PyOpenCV module **(B)**. Linear (Gaussian) filter followed by edge enhancement (sharpening) or non-linear filter (median filter) are suitable in different species based on their cellular features (see text). A scan line of the Golgi apparatus drawn with Fiji in *Emiliania huxleyi* shows the impact of different filters on the profile plot of the flattened membrane-enclosed disks (the cisternae). Red: original image. Blue: A Gaussian filter smooths the edges and some membranes disappear. Purple: sharpening after application of the Gaussian filter allows recovery of some image details after smoothing edges. Green: The median filter is less sensitive to edges. Image processing was done with 3D Slicer for segmentation **(C)**, MeshLab and Paraview, for editing and visualisation, respectively **(D)**. The STL and Trimesh python packages were used to quantify volumes, surfaces and distances **(E)**. From a watertight mesh, surface is obtained by summing the surface of each individual triangle present in the mesh. Volume is computed with a volume integral formula discretized over tetrahedrons as obtained from (Zhang and Chen, 2001). Distances between nearby point clouds of different objects within a cell were used to estimate organelle proximity. The whole process requires 10 to 15 days for a microalgal cell with a cell diameter comprised between 2 μm to 8 μm.

Segmentation of organelles, vesicular networks, vacuoles and storage compartments was carried out with 3D Slicer software (Kikinis et al., 2014) (www.slicer.org, Figure 1C), using a manually-curated, semi-automatic pixel clustering mode (3 to 10 slices are segmented simultaneously for a given Region Of Interest, (ROI)). This method is optimal in terms of image processing speed and allowed fast and accurate identification of subcellular structures when compared to a fully manual segmentation while avoiding possible artefacts (e.g. the presence of misclassified pixels). The segmented stacks, represented in memory as simple binary masks, were converted into 3D models using the “Model maker” module from 3D Slicer (Figure 1C), and then edited with the MeshLab software (Cignoni et al., 2008; Ranzuglia et al., 2012) to eliminate ‘isolated islands’, which were erroneously annotated as ROIs. We also used MeshLab to simplify meshes (‘mesh decimation’, Figure 1D), i.e. to reduce the model nodes and faces down to 25% of the original data without modifying morphometric values, such as surfaces and volumes (Supplementary Table 2). Every 3D model was imported into Paraview (Figure 1D) (Ahrens et al., 2005) to visualise 3D objects and understand their relationship. Blender (www.blender.org) was used for object animation (Supplementary Video 7). Morphometric analyses were performed using the Python module numpy-slt (https://github.com/WoLpH/numpy-stl), which provided estimates very similar to those obtained with the geometry tool of MeshLab and 3D Slicer (Supplementary Table 3). This Python package is faster than MeshLab, with obvious advantages in terms of analysis of large files (> 500 MB). Surfaces and volumes were computed using the discrete mesh geometry, surface being computed directly from mesh triangles, and volume being obtained from the signed volume of individual tetrahedrons, assuming a closed surface (watertight mesh Figure 1E). Using the Trimesh Python module, the minimal distance between two meshes was calculated based on the closest points between two triangular meshes. Hence, the surface area contacts were quantified based on: *i*) calculating the minimal distance between each vertex of the plastid mesh to the mitochondria mesh (for 3 cells of every species), and then by *ii)* gathering mesh vertices according to a given distance threshold to generate contact surface. A distance threshold ≤ 30 nm was chosen as representative of an interaction between nearby organelles, on the basis of previously established morphometric analysis in animal and plant cells (Helle et al., 2013; Mueller-Schuessele and Michaud, 2018; Scorrano et al., 2019).

Overall, this entirely free-access analytical pipeline meets all the quality criteria for selecting, filtering and classifying ROIs and for building optimal 3D structure models for morphometric analysis. It allows us to generate a complete cell model (from cryofixation to 3D visualization and morphometric analysis) in 10-15 days.

### 2 Cellular architectures of phytoplankton

Our FIB-SEM tomograms highlighted different cell architectures in the studied microalgae (e.g. coccolithophores in *Emiliania*, the raphe in *Phaeodactylum*, Micromonas’ flagellum), reflecting their different phylogenetic origin (Figure 2), but also showed several common topological characteristics. Internal cell structures (Figure 3B) include: *i*) organelles (nucleus-blue, plastid-green and mitochondria-red), *ii*) the cytosol plus internal vesicular networks (grey): Golgi apparatus, vacuoles and storage compartments. The latter group includes Ca^2+^-rich storage bodies and forming coccolith in *Emiliania* (Gal et al., 2018; Sviben et al., 2016), carbon-rich structures in *Pelagomonas* (Andersen et al., 1993), lipid droplets in *Phaeodactylum* (Lupette et al., 2019), starch sheath surrounding the pyrenoid in *Micromonas* (Lopes Dos Santos et al., 2017) and *Symbiodinium* and vacuoles of different sizes in *Phaeodactylum*, *Galdieria* and *Micromonas* (the so-called impregnated bodies, (Lopes Dos Santos et al., 2017)).

**Figure 2:**
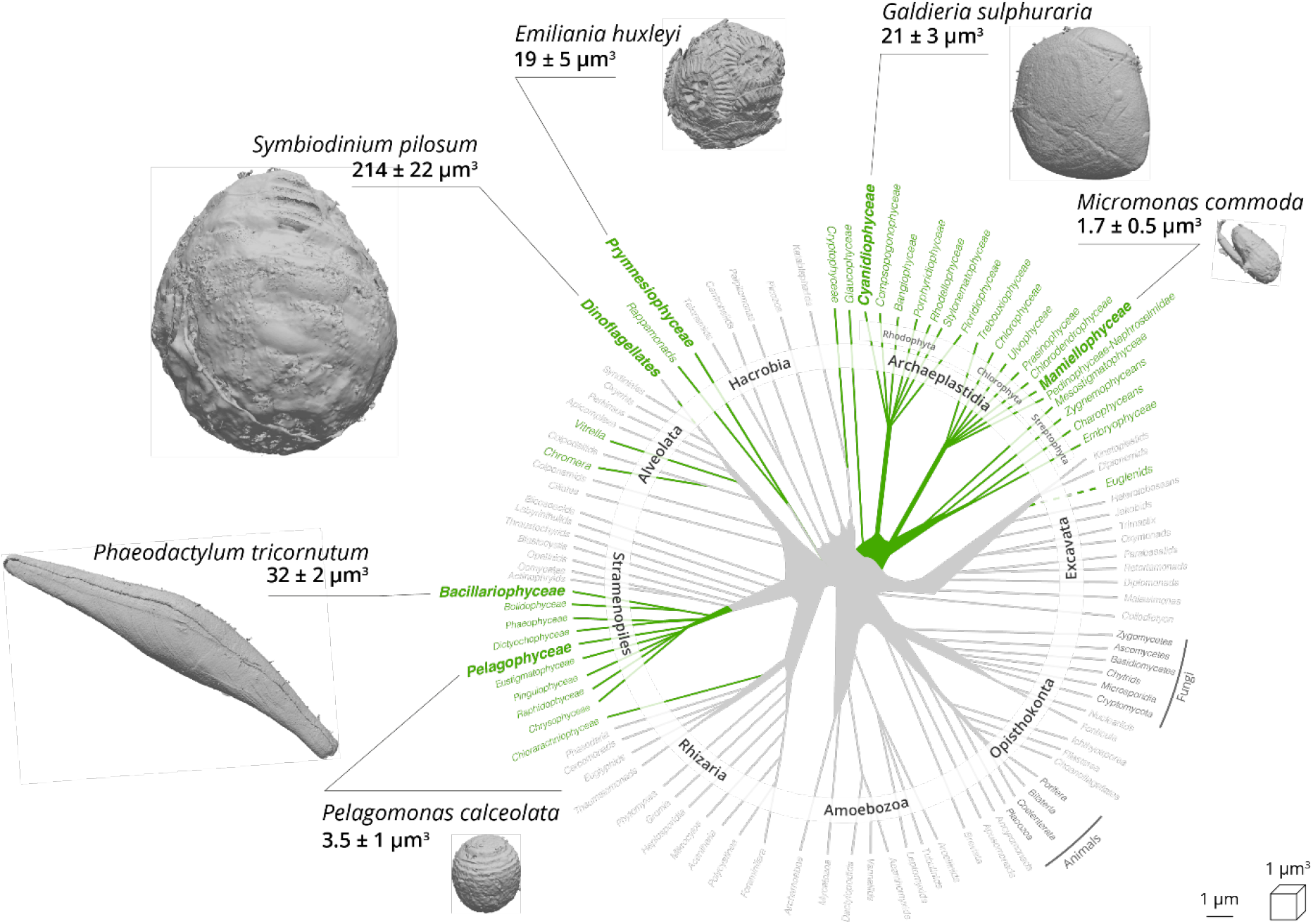
External feature of selected phytoplankton cells revealed by FIB-SEM imaging. Green lineages of the phylogenetic tree of eukaryotes represent photosynthetic eukaryotes (adapted from (Decelle et al., 2015)). A 3D scan view of cell morphology of selected phytoplankton members (Mammiellophyceae: *Micromonas commoda,* Pelagophyceae: *Pelagomonas calceolata;* Prymnesiophyceae: *Emiliania huxleyi;* Cyanidiophyceae: *Galdieria sulphuraria;* Bacillariophyceae: *Phaeodactylum tricornutum;* Dinophyceae: *Symbiodinium pilosum*) is shown with a linear scale bar of 1μm and a voxel scale of 1 μm^3^. Specific cellular features (cell walls, the flagellum in *Micromonas*, coccolithophores in *Emiliania*, the raphe in *Phaeodactylum*) are visible. Three cells were reconstructed and their morphometry measured for every phytoplankton taxa.

**Figure 3:**
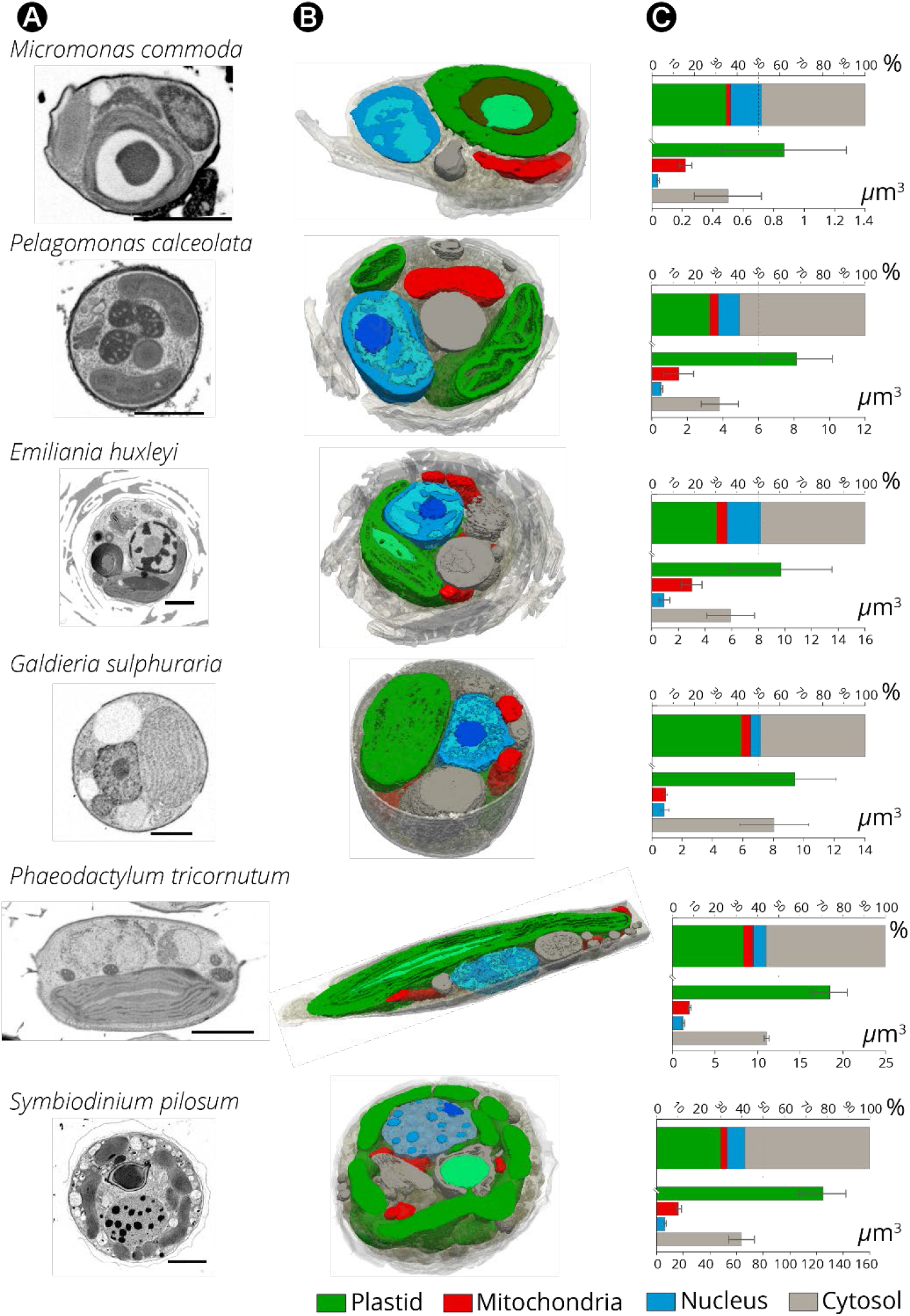
Internal cell architecture of phytoplankton cells as revealed by FIB-SEM imaging. **(A):** 2D FIB-SEM frames of every phytoplankton species. Scale bar: 2 μm **(B):** Sections of cellular 3D volumes, based on FIB-SEM imaging of whole cells of *Micromonas* (stack of frames in **Supplementary Video 1**), *Pelagomonas* (**Supplementary Video 2**), *Emiliania* (**Supplementary Video 3**), *Galdieria* (**Supplementary Video 4**) *Phaeodactylum* (**Supplementary Video 5**) and *Symbiodinium* (**Supplementary Video 6**). Images highlight the main subcellular compartments: green: plastids (containing thylakoids and pyrenoids - light green-in some cell types); red: mitochondria; blue: nuclei (containing euchromatin –light blue-heterochromatin - blue- and the nucleolus - dark blue-); grey: the Golgi apparatus vacuoles and storage compartments and the cytosol. **(C):** Volume occupancy by the different subcellular compartments in different microalgal cells. Top plot: % of occupation; bottom plot: absolute volume sizes (N=3 ± SD)

The plastids were cup-shaped in *Galdieria*, *Pelagomonas*, *Emiliania*, lobed in *Symbiodinium* (Blank, 1987), globular in *Micromonas* and elongated in *Phaeodactylum* (Figure 3B and Figure 4A). When distinguishable, photosynthetic membranes (thylakoids) were organised in layers of a few stacks, without the typical differentiation in grana and stromal lamellae observed in vascular plants and green algae (Mustardy and Garab, 2003). Mitochondria were characterized by extremely variable shapes between species and within cells of the same species (e.g. Supplementary Figure 1 in the case of *Emiliania*), reflecting the dynamic nature of this organelle. Transient and rapid morphological changes of the mitochondria through cycles of fission and fusion are probably crucial for phytoplankton acclimation, as previously concluded for plant and animal cells (Tilokani et al., 2018). Tilokani et al., 2018). The nucleus had a more consistent shape and was closely associated to the plastid in most cases, via the fourth envelope membrane in secondary plastids (i.e. *Phaeodactylum* (Flori et al., 2016)).

**Figure 4:**
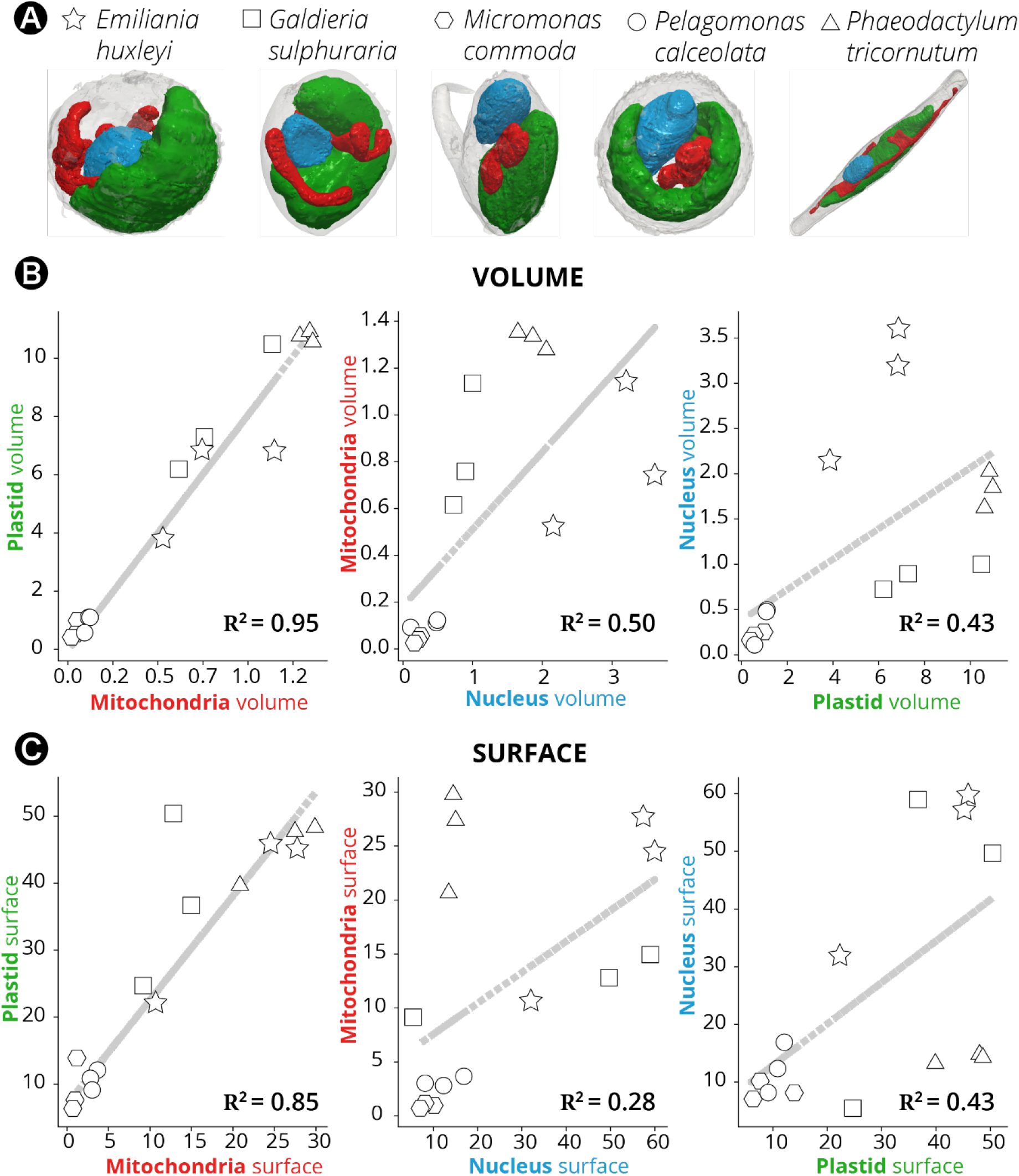
Morphometric analysis of phytoplankton members. **(A):** 3D topology of the main organelles (green: plastids; red: mitochondria; blue nuclei) in the different cell types. **(B):** Volume relationships in different subcellular compartments, as derived from quantitative analysis of microalgal 3D models. **(C):** Surface relationships in different subcellular compartments, as derived from quantitative analysis of microalgal 3D models. Three cells were considered for every taxum. hexagons: *Micromonas*; circles: *Pelagomonas*; stars: *Emiliania*; squares: *Galdieria*; triangles: *Phaeodactylum*. *Symbiodinium* cells were not considered in this analysis, because their size, which largely exceeds the other (**Supplementary Figure 2**), prevents a correct analysis of the volume/surface relationships.

The largest organelle was the plastid, which occupied 30 to 40% of the cell volume depending on the microalga (Figure 3C). The nucleus occupied 5 to 15% of the cell volume while the mitochondria occupied a lower cell volume (2 to 5%, see also Supplementary Table 4). Altogether, the organelles (nuclei, plastids and mitochondria) filled a relatively constant fraction (40 to 55 %) of the total cell volume in the different lineages studied. Networks of internal vesicles, the Golgi apparatus, vacuoles and storage compartments (e.g. lipid droplets, starch granules, nutrient storage, etc) and the cytosol occupied the other half with a high variability in terms of volume occupancy. We interpret this conservation of organelle volumes and the variability of the other compartments as the signature of evolutionary constraints that preserve essential cellular functions (gene expression, energy production, anabolism/catabolism), while leaving metabolic flexibility to allow the storage of assimilated nutrients, particularly carbon.

Thanks to the possibility offered by our approach to perform quantitative surface and volumetric analyses, we sought possible relationships between the organelles (Figure 4A) in the different taxa, to reveal evolutionary preserved morphological characteristics. This analysis was initially biased by the presence of *Symbiodinium* (Supplementary Figure 2). These dinoflagellate cells, being much larger than the others, result in the clustering of data into two cells group (*Symbiodinium* on the one side and all the other cells on the other one) leading to the observation of apparent linear relationships between all the parameters considered. Excluding *Symbiodinium* from the analysis removed this bias and unveiled the existence of a tight correlation between plastids and mitochondria in terms of their volume (the Pearson correlation coefficient, R, being 0.95, Figure 4B) and surface area ratios (R = 0.85, Figure 4C). This conserved topological feature corroborates the previous molecular and physiological evidence in diatoms (Bailleul et al., 2015; Kim et al., 2016) for energetic interactions between plastids and mitochondria. Conversely, no significant correlation was found between the volume/surface ratio of the nucleus and the mitochondria or plastid (R ≤ 0.5).

Plastid-mitochondria interactions may rely on physical contacts, as already proposed (Flori et al., 2017; Mueller-Schuessele and Michaud, 2018). We explored this hypothesis by quantifying the contact areas (defined as a minimal distance of < 30 nm (Scorrano et al., 2019)) between plastids and mitochondria in the different species. We detected areas of contact between the two organelles in all microalgae (Figure 5), further corroborating the idea of specific and conserved interactions between these two cellular engines. However, different extents and distributions of surface area contacts were found in the different species (Figure 5). While approximately 5% of the total surface of the plastid was interacting with mitochondria in the diatom *Phaeodactylum*, this interaction was reduced to 1-2% of the plastid surface in *Micromonas, Emiliania* and *Galdieria* and to less than 1 % in *Pelagomonas* and *Symbiodinium*. Thus, although plastid-mitochondrial connections are observed in unicellular phytoplankton eukaryotes (Figure 4), their extent varies depending on the species (Figure 5), suggesting that the whole process could be dynamic throughout the life cycle and environmental conditions, possibly being mediated by species-specific processes.

**Figure 5:**
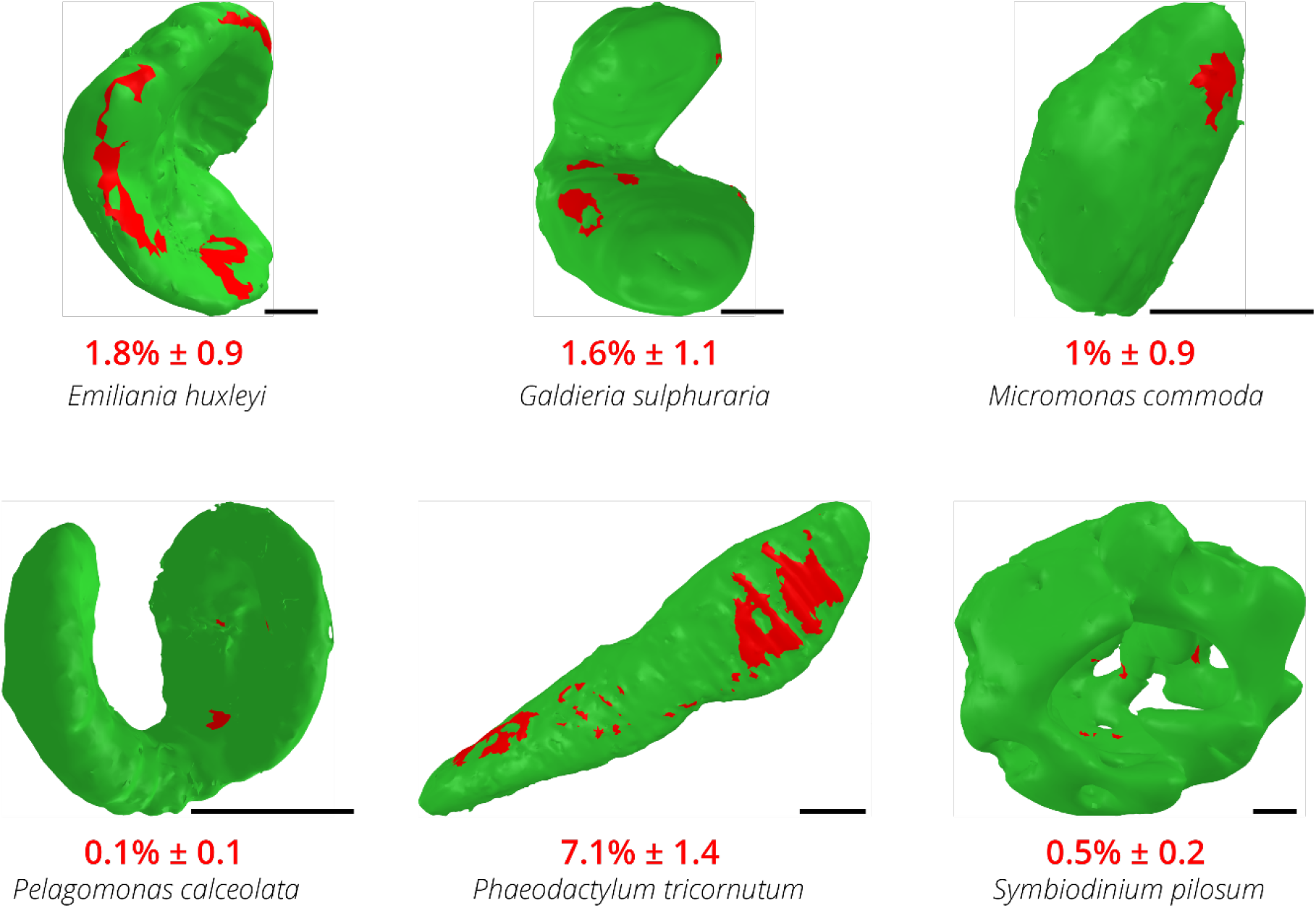
Contact surface areas between plastids and mitochondria in different phytoplankton members. Green: plastid surface. Red spots: contact points (i.e. points at a distance ≤ 30 nm) between mitochondria and plastids. Scale bars: 1 μm.

### 3 Subcellular features

Besides providing information on the topologies of whole cells, our 3D images had enough resolution to explore the sub-organelles features of the different microalgae (Figure 6, Supplementary Figure 3). The plastid volumes were mainly occupied by thylakoid membranes and, when present, by the pyrenoid, the compartment containing Rubisco (Figure 6A). This compartment occupied between 2-9% of the plastid volume (Figure 6B). In two species (*Phaeodactylum* and *Emiliania*), we observed thylakoids crossing the pyrenoid matrix (Figure 6A). However, these pyrenoid membranes (also called pyrenoid tubules in *Chlamydomonas reinhardtii*, (Engel et al., 2015)) exhibited very different topologies: we observed parallel stacks in the diatom and a more branched structure in *Emiliania*, reminiscent of that recently reported in *Chlamydomonas reinhardtii* (Engel et al., 2015; Meyer et al., 2016). Our FIB-SEM images showed that *Micromonas* and *Symbiodinium* contained thylakoid-free pyrenoids, that were almost completely surrounded by starch sheaths (Figure 6A). Few stalks ensure the connection between pyrenoid and the plastid, possibly to facilitate the diffusion of Rubisco substrates and products as previously proposed (Badger and Price, 1994; Engel et al., 2015; Moroney and Mason, 1991), see also the review by (Meyer et al., 2017). Unlike *Micromonas*, the pyrenoid of the dinoflagellate *Symbiodinium* was not centred in the plastid, but instead protruded towards the cytosol, being surrounded by a shell of cytosolic rather than stromal starch (Dauvillee et al., 2009; Meyer et al., 2017; Van Thinh et al., 1986).

**Figure 6.**
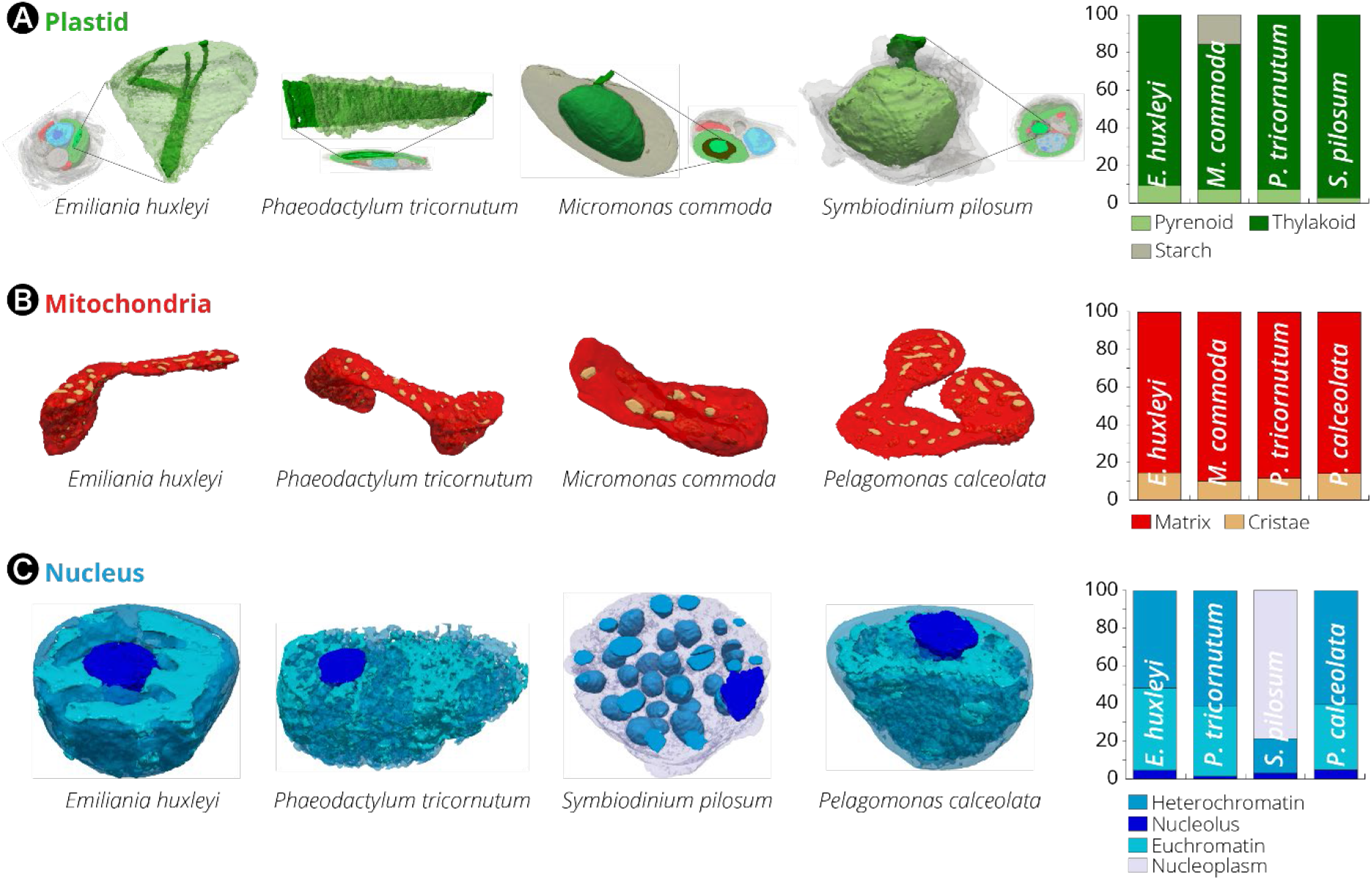
Subcellular features of different phytoplankton taxa. **(A):** The CO_2_-fixing compartment (pyrenoid) topology in *Phaeodactylum, Emiliania, Micromonas and Symbiodinium* cells. The 3D reconstruction displays the thylakoid network (dark green) crossing the pyrenoid matrix (light green). If present (*Micromonas* and *Symbiodinium*), a starch layer surrounding the pyrenoid is shown in grey. The histogram recapitulates volume occupancy by sub-plastidial structures (thylakoids, matrix, starch). Note that starch is cytosolic in *Symbiodinium*, and therefore its volume is not considered in the chart. **(B):** Mitochondrial features. Topology of mitochondrial compartments. Red: mitochondrial matrix; yellow: cristae. The histogram recapitulates volume occupancy by mitochondrial sub-compartments (in the matrix and within the cristae). Despite changes in the mitochondria morphology, likely reflecting the dynamic character of these organelles (**Supplementary Figure 1**), the ratio between the subcellular compartment volumes is relatively constant in the different cells. **(C):** Nuclear features. Topology of the nucleus. Light blue: euchromatin; blue: heterochromatin. Nucleoids (dark blue) are also visible. The histogram recapitulates volume occupancy by the different types of chromatin. Different levels of DNA condensation are visible. DNA is present in the form of compact chromosomes in *Symbiodinium*, possibly leaving a fraction of the nucleoplasm without chromatin (grey).

This close relationship between starch and pyrenoids, and the fact that the starch sheath around the pyrenoid is rapidly formed in response to a decrease in the CO_2_ concentration (Ramazanov et al., 1994) indicate a possible role of the starch sheath in the Carbon Concentrating Mechanism (CCM) of microalgae. One possible function (limiting CO_2_ diffusion out of the pyrenoid and away from Rubisco) was challenged by the characterisation of microalgal mutants devoid of starch (del Pino Plumed et al., 1996; Villarejo et al., 1996, see however (Toyokawa et al., 2020)), leading instead to the hypothesis of a positive role of the pyrenoid in the local deposition of starch (review in (Meyer et al., 2017)). In this context, one could speculate that the proximity between the reserve polymers and the pyrenoid reflects the fact that the first step in starch biosynthesis (the formation of ADP-glucose by ADP-glucose pyrophosphorylase) is stimulated by 3PGA (3-phosphoglyceric acid) - a direct product of CO_2_-fixing Rubisco during the Calvin Benson cycle. Thus, starch would be synthesized where the concentration of 3PGA is the highest, i.e. near Rubisco in the pyrenoid. However, it is unclear if this explanation would hold in *Symbiodinium*, the substrate for which may be UDP-glucose rather than ADP-glucose (Viola et al., 2001)). More recently, the starch sheath was proposed to help limiting the number of pyrenoids that form within the plastid (Itakura et al., 2019).

Despite the differences in the pyrenoid topology, the ratio of pyrenoid/plastid volumes was preserved in three out of the four microalgae lineages where this compartment was present (7%, 9%, 7% for *Phaeodactylum*, *Emiliania*, *Micromonas*, respectively, Figure 6A and Supplementary Table 4). This constant ratio highlights the importance of maintaining a proper balance between the subcompartments producing light-dependent energy (the photosynthetic membranes) and the light-independent one (the CO_2_ fixing compartment). An exception to this observation is *Symbiodinium*, where the pyrenoid occupies a much lower fraction of the plastid volume (2% ca). Our quantitative morphometric analysis provides a rationale for this exception. We found that the pyrenoid surface/volume ratio (an important parameter for gas exchange in this compartment, and therefore for CO_2_ assimilation) is 15-20 in *Phaeodactylum*, *Emiliania*, *Micromonas* (Supplementary Table 4) but only around 5 in the dinoflagellate. This possibly explains why the large increase in the plastid volume of *Symbiodinium* (60 μm^3^ vs 10 μm^3^ in *Phaeodactylum*, 6 μm^3^ in *Emiliania* and 0.5 μm^3^ in *Micromonas*, Supplementary Table 4) is not paralleled by a commensurate expansion of the pyrenoid volume (1.8 vs 0.8, 0.6 and 0.05 μm^3^ respectively, in the four algae). A much lower surface to volume ratio may represent a functional constraint for carbon assimilation.

Overall, the volumetric analysis of the pyrenoid suggests that both the surface to volume ratio and the volumetric ratio between the plastid and the pyrenoid are important parameters for the photosynthetic metabolism. This concept of constant volumetric ratios within energy producing organelles was also found at the level of mitochondria. In this case, we found that the ratio between the volume inside the cristae and the matrix (Figure 6B) is relatively constant in all cells (12%, 14%, 15%, 10% in *Phaeodactylum*, *Pelagomonas*, *Emiliania* and *Micromonas*, respectively, Figure 6B), despite differences in the shape (Fig 4A) and overall volumes of their mitochondria (Figure 3B supplementary Figure 1).

Our 3D analysis allows us to reveal some species-specific subcellular features. In the nucleus, we observed differences in the extent of DNA condensation (Figure 6C), possibly reflecting differences in the transcription activity of the different cultures at the time they were fixed. Patterns vary from the largely euchromatic nucleus of *Phaeodactylum* (60% of the volume occupied by non-condensed DNA) to the fully compacted dinokaryon in *Symbiodinium*, where around 100 chromosomes of different sizes were clearly distinguishable (Supplementary Figure 4).

Finally, we could reconstitute in 3D most of the known steps of the process of biomineralization, known as coccolithogenesis (for reviews see (de Vargas et al., 2015; Taylor et al., 2017)) within a single cell (Figure 7A). This process starts with the formation of a proccolith ring within Golgi-associated vesicles (during the nucleation phase, Taylor et al., 2017 see also Sviben et al., 2016; Gal et al., 2018), which were clearly visible inside the cell (Figure 7B). Coccoliths become more structured insofar as the vesicles detached from the Golgi apparatus, in a step called maturation (Taylor et al., 2017). Typical features of this phase (prococcolith rings, the organic base plate scale-OBPS-that allows nucleating the CaCO3 crystals and, possibly, the reticular body (Beuvier et al., 2019)) are visible in our 3D images (Figure 7B). Coccolith are finally secreted to the cell surface (during the secretion phase (Taylor et al., 2017), where they constitute the inner layer the “coccosphere”, (Figure 7C, N°1) i.e. the cell exoskeleton. Outside the cell, coccoliths progressively move towards the outermost part of the (Figure 7C, N°2 to N°5). Therefore, the most recently produced coccoliths (N° 1) lie beneath older ones (N° 5).

**Figure 7.**
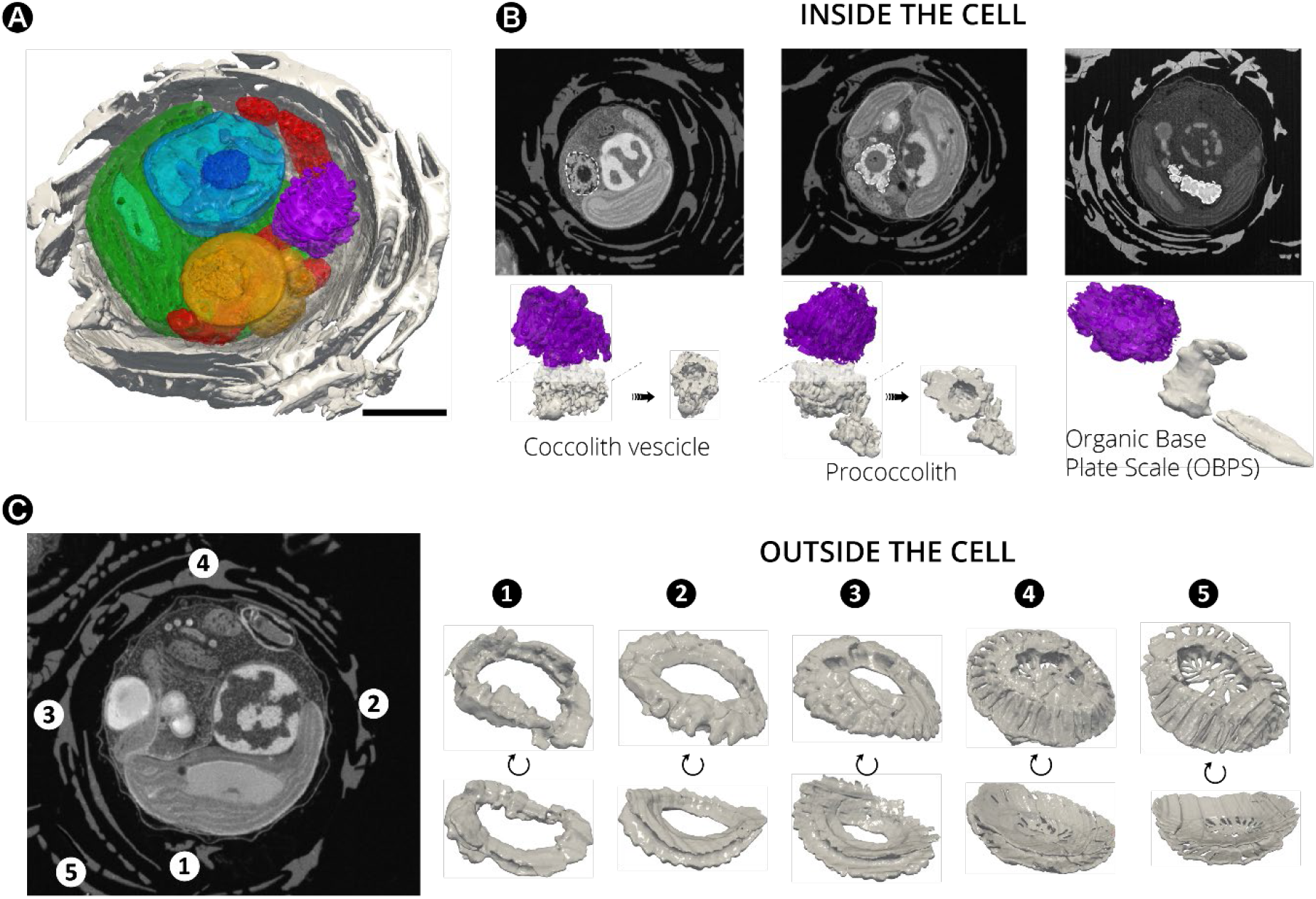
Morphogenesis of Coccolithophores in Emiliania huxleyi. **(A):** Sections of cellular 3D volume. **(B):** FIB-SEM sections and 3D rendering of *Emiliania* structures showing changes in the morphology of nucleating coccoliths (grey) during coccolithogenesis. The process starts with the ‘nucleation’ phase (Taylor et al., 2017), where coccolith vesicles are formed in close contact to the Golgi apparatus (purple) and the storage compartments (yellow). Within the cell, typical elements of the ‘maturation’ phase (prococcoliths) are also visible. During the ‘secretion’ phase, coccoliths are released outside the cells making the Organic Base Plate Scale (OBOS) easy detectable in the cell. Secreted coccolith are deposited **(C)** in the inner part of the coccosphere (e.g. N° 1 and N° 2). This process triggers the movement of more mature coccoliths (e.g. N°3, N°4 and N°5) towards the outermost part of the exoskeleton. Therefore, the most recently produced coccoliths (N° 1) lie beneath older ones (N° 5). Note that coccoliths in the inner layers of the coccosphere have still partially incomplete structural features. Scale bar: 1 μm. Additional morphological features of *Emiliania huxleyi* are visible in the **Supplementary Video 7.**

## Conclusions

When compared to other imaging approaches based on electron microscopy (such as cryo-electron tomography in a TEM), the FIB-SEM approach reported here has the clear advantage of providing a contextual 3D view of whole cells, of very different sizes, at high-resolution. In principle, different types of cells can be mixed before cryo-preparation of the sample, allowing a fast and reliable comparison for environmental microbiology. The relative slowness of FIB-SEM acquisitions (1 or 2 day per sample) can be mitigated by the automated acquisition of image stacks for several cells in parallel. The pipeline presented here, entirely based on open access software, improves the speed and reliability of image analysis and 3D reconstruction. This approach can be applied to samples from different environments and can be used for quantitative comparative analysis of different species to reveal important links between cell structures and physiological/metabolic responses. Here, we focused on a limited set of cell types and only one growth condition. However, many conditions can potentially be compared to visualize the acclimation strategies of photosynthetic microbes to their environment, filling the knowledge gaps between physiology, cell biology, ecosystem structure and function. This could be relevant for predicting the consequences of climate change (Winder and Sommer, 2012), which will likely affect the size and morphology of phytoplankton as temperature, acidity and the availability of nutrients change (Intergovernmental Panel on Climate, 2014). We have demonstrated that, thanks to the nanometric resolution of SEM, morphological analysis can be extended to subcellular and even sub-organellar structures, and can potentially help understanding physiological responses at the macromolecular level. This is illustrated by the finding of conserved relationships between plastidial and mitochondrial volumes, possibly to maintain optimal signalling and energy flow during the assimilation of carbon (Bailleul et al., 2015). Our observation of different intermediates of biomineralisation, a fundamental biogeochemical process that initiates carbon deposition in the oceans is another nice example. The major challenges in the future will be to correlate this approach with cryo-electron tomography in order to improve the resolution of specific subcellular structures to the molecular level, and to combine it with chemical imaging (Decelle et al., 2020) and fluorescence measurements with fluorophores/antibodies, in order to allow correlative microscopic studies on phytoplankton (Sartori et al., 2007; Stephens and Allan, 2003).

## Material and methods

### Species

The species used in this work (Supplementary Table 1) were chosen on the basis of two criteria: i) they must be representative of phytoplankton taxa that are ecologically relevant and *ii*) they can be grown in laboratory conditions.

### Algal cultivation

*Phaeodactylum tricornutum* Pt1 strain (CCAP 1055/3) was obtained from the Culture Collection of Algae and Protozoa, Scottish Marine institute, UK. Cells were grown in the ESAW (Enriched Seawater Artificial Water) medium (Berges et al., 2004) in 50 mL flasks in a growth cabinet (Minitron, Infors HT, Switzerland), at 19°C, a light intensity of 40 μmol photon m^−2^s^−1^, a 12-h light /12-h dark photoperiod and shaking at 100 rpm. *Galdieria sulphuraria* SAG21.92 was obtained from the University of Dusseldorf (Germany) and was grown in sterile 2XGS modified Allen medium, pH 2.0 (Allen, 1959) at 42°C under the same light conditions. Cells were grown in 250 mL flasks (50 mL culture volume). *Micromonas commoda* RCC 827, *Pelagomonas calceolata* RCC 100, *Emiliania huxleyi* RCC 909 in K medium at 20°C, and *Symbiodinium pilosum* RCC 4014 in F/2 medium at 20°C were obtained from the Roscoff Culture Collection (Vaulot et al., 2004) and maintained in the same medium and temperature condition without agitation. Cells were kept at a light intensity of 60-80 μmol photon m^−2^s^−1^, a 12-h light /12-h dark photoperiod, without shaking.

### Sample preparation for electron microscopy

Sample preparation protocols were adapted from (Decelle et al., 2019) to optimize the contrast for 3D electron microscopy imaging and therefore facilitate image segmentation through pixel classification. Live cells were cryofixed using high-pressure freezing (HPM100, Leica) in which cells were subjected to a pressure of 210 MPa at −196°C, followed by freeze-substitution (EM ASF2, Leica). Prior to cryo-fixation, the microalgal cultures were concentrated by gentle centrifugation for 10 min (800 rcf). For the freeze substitution (FS), a mixture 2% osmium tetroxide and 0.5% uranyl acetate in dried acetone was used. The freeze-substitution machine was programmed as follows: 60-80 h at −90°C, heating rate of 2°C h^−1^ to −60°C (15 h), 10-12 h at −60°C, heating rate of 2°C h^−1^ to −30°C (15 h), and 10-12 h at −30°C, quickly heated to 0°C for 1 h to enhance the staining efficiency of osmium tetroxide and uranyl acetate and then back at −30°C. The cells were then washed four times in anhydrous acetone for 15 min each at −30°C and gradually embedded in anhydrous araldite resin. A graded resin/acetone (v/v) series was used (30, 50 and 70% resin) with each step lasting 2 h at increased temperature: 30% resin/acetone bath from −30 °C to −10 °C, 50% resin/acetone bath from −10 °C to 10 °C, 70% resin/acetone bath from 10 °C to 20 °C. Samples were then placed in 100% resin for 8-10 h and in 100% resin with the accelerator BDMA for 8 h at room temperature. Resin polymerization finally occurred at 65 °C for 48 h.

### FIB-SEM acquisition imaging

Focused ion beam (FIB) tomography was performed with either a Zeiss NVision 40 or a Zeiss CrossBeam 550 microscope, both equipped with Fibics Atlas 3D software for tomography. The resin block containing the cells was fixed on a stub with carbon paste, and surface-abraded with a diamond knife in a microtome to obtain a perfectly flat and clean surface. The entire sample was metallized with 4 nm of platinum to avoid charging during the observations. Inside the FIB, a second platinum layer (1-2 μm) was deposited locally on the area analysed. The sample is then abraded slice by slice with the Ga^+^ ion beam (generally with a current of 700 nA at 30 kV). Each freshly exposed surface is imaged by scanning electron microscopy (SEM) at 1.5 kV and with a current of ~ 1 nA using the in-lens EsB backscatter detector. For algae, we generally used the simultaneous milling and imaging mode for better stability, and with an hourly automatic correction of focus and astigmatism. For each slice, a thickness of 8 nm was removed, and the SEM images were recorded with a pixel size of 8 nm, providing an isotropic voxel size of 8×8×8 nm^3^ Whole volumes were imaged with 800 to 1000 frames, depending on the species. Due to its reduced cell dimensions, the voxel size was reduced to 4×4×4 nm^3^ in the case of *Micromonas*, resulting in higher resolution datasets with approximately 350-500 frames/cell.

### Image processing

As a first step of image processing, ROIs containing cells were cropped from the full image stack. This was followed by image registration (stack alignment), noise reduction, semi-automatic segmentation of ROIs, 3D reconstruction of microalgae cells and morphometric analysis. Several problems may be encountered during these steps. Raw stacks consist of big data (50 to 100 GB for the whole imaged volume, containing several cells) that do not necessarily fit into the computer main memory (RAM). Moreover, cryo-substituted cells generate less contrasted images than cells prepared with chemical fixation. Therefore, the first step in building a robust 3D model consists in ‘isolating’ a given ROI (e.g. an organelle) from other compartments, to obtain a smaller stack size that can be easily worked with (in practice, we worked with substacks that were around 10% of the original stack size).

Single cells were isolated by cropping in 3-dimensions using the open software Fiji (https://imagej.net/Fiji). Image misalignment was corrected using template matching (“align slices in stack“) option implemented in Fiji. This function tries to find the most similar image pattern in every slice and translates them to align the landmark pattern across the stack (https://sites.google.com/site/qingzongtseng/template-matching-ij-plugin) (Videos S1-S6). Aligned image stacks were filtered to remove noise using Python (Oliphant, 2007) and OpenCV (OpenCV. 2015, Open Source Computer Vision Library programming tools). Filtering techniques were chosen to highlight contours while removing noise in the images. Depending on the species, we found that the osmium staining was not homogeneously distributed. Therefore, it was not possible to filter raw datasets of every species with the same method. Application of a Gaussian filter followed by sharpening to remove noise and enhance contours, which is widely used and easy to implement (Russo, 2002), was used to process raw datasets of *Emiliania*, *Micromonas*, *Phaeodactylum* and *Pelagomonas*. However, this method was less effective when applied to raw datasets of *Galdieria* and *Symbiodinium*, where using the median filter turned out to be a better de-noising option.

### Segmentation

The filtered stacks were loaded into 3D Slicer to be processed. By selecting the model editor, we ‘coloured’ the ROIs using paint tools and adjusted the threshold range for image intensity values. The ROIs were annotated and the corresponding label map was run into the model maker module, to generate corresponding 3D models that were exported in different formats (.stl, obj, vtk, ply,.mtl). For further analysis, we used the .stl mesh, which proved to be more suitable for 3D analysis in our workflow (Supplementary Table 2).

### 3D reconstructed model

A 3D filtering process was needed to clean the model and reduce the size of the file (see Supplementary Table 2). In our case, 3D models generated by 3D Slicer were imported into the open source software MeshLab (Cignoni et al., 2008) to clean the model by automatically removing isolated islands. We also performed a remeshing process to facilitate 3D modelling, visualization and animation. The edited model was then imported into Paraview software to capture a 2D representation of the 3D reconstruction. We used Blender for model animation (see an example in Supplementary Video 7).

### Morphometric evaluations

Measurement of volumes, surface area, and the minimum distance between meshes) were performed using Numpy-STL (https://pypi.org/project/numpy-stl/) and TRIMESH (https://trimsh.org/trimesh.html) packages of Python (Supplementary Table 3).

### Surface and volume measurements

To compute the surface, we iterated over all the triangles of the mesh. The computation of the cross product between two edges of a given triangle gives a vector whose magnitude is twice the area of said triangle. Then, the sum of all these areas provides the total surface area of the mesh. We then computed the signed volume of all tetrahedrons, which goes from the origin (0,0,0) to each triangle present in the mesh. Assuming a closed surface (watertight mesh), summing all those volumes give the volume of the mesh (Zhang and Chen, 2001). A simple implementation of those algorithms is provided by the authors at https://gitlab.com/clariaddy/stl_statistics.

### Distance between organelles

Given two meshes, we computed the pointwise distance from the first mesh to the second one. For each vertex in the first mesh, we computed the minimal distance to the second mesh using the Trimesh Python module (https://github.com/mikedh/trimesh). Using this method, every vertex of the mitochondria model was compared to all vertices contained in the plastid model to calculate the intermesh distance. After having set a biological meaningful threshold distance (≤ 30 nm, (Helle et al., 2013, Scorrano et al., 2019)), we generated subsets of plastid-mitochondria points that meet this criterion and reconstructed the matching surface using face data. The corresponding surfaces were then compared to the total plastid surface (https://gitlab.com/clariaddy/mindist).

## Acknowledgements

This project received funding from the European Research Council: ERC Chloro-mito (grant no. 833184). Research was also supported by a Défi X-Life grant from CNRS, funds from the CEA DRF impulsion FIB-Bio program, the LabEx GRAL (ANR-10-LABX-49-01), financed within the University Grenoble Alpes graduate school (Ecoles Universitaires de Recherche) CBH-EUR-GS (ANR-17-EURE-0003) and the ANR ‘Momix’ (Projet-ANR-17-CE05-0029). This project also received funds from the European Union’s Horizon 2020 research and innovation programme CORBEL under the grant agreement No 654248. This work used the platforms of the Grenoble Instruct-ERIC centre (ISBG; UMS 3518 CNRS-CEA-UGA-EMBL) within the Grenoble Partnership for Structural Biology (PSB), supported by FRISBI (ANR-10-INBS-05-02) and GRAL. The electron microscope facility is supported by the Auvergne-Rhône-Alpes Region, the Fondation Recherche Medicale (FRM), the funds FEDER and the GIS-Infrastructures en Biologie Santé et Agronomie (IBISA). J.D. was supported by ATIP-Avenir program. C.U. is supported by a joint UGA-ETH Zurich PhD grant in the framework of the “Investissements d’avenir” programme (ANR-15-IDEX-02). The authors thank the Roscoff Culture Collection that provided phytoplankton strains and Noan Le Bescot (Ternog Design) for help in the conception and realisation of the figures of this article.

---

**Supplementary Figure 1:**
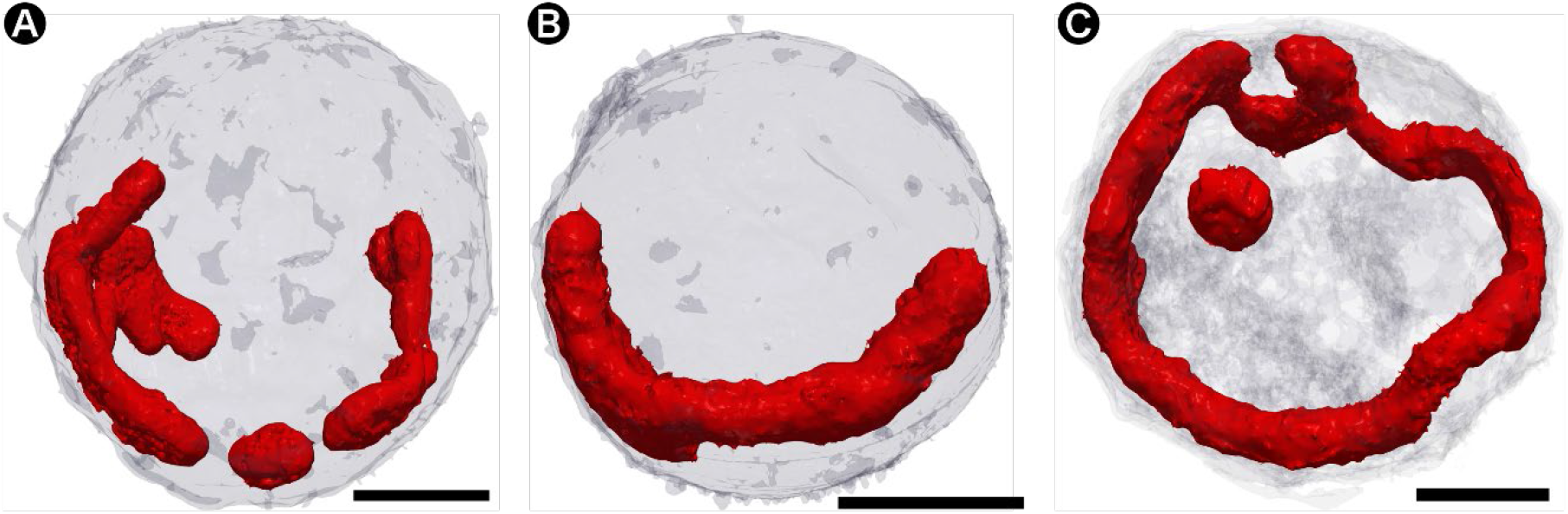
The different morphologies of mitochondria in three cells (a to c) of Emiliania huxleyi (Prymnesiophyceae) The different 3D topology of the mitochondria (red) in *Emiliania* (light grey) highlights the dynamic character of this organelle. Scale bar: 1 μm.

**Supplementary Figure 2.**
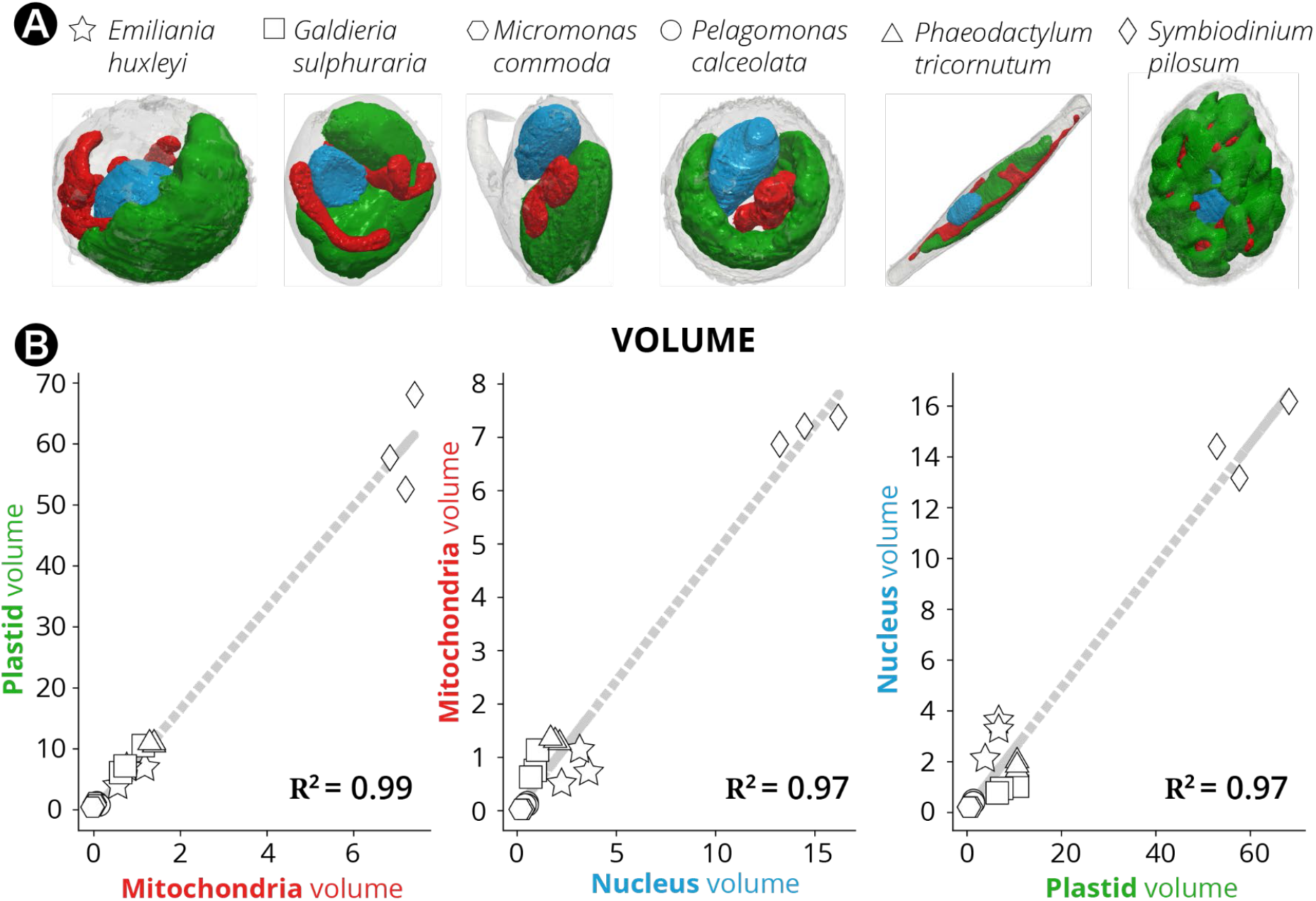
Volumes and surfaces relationship in different subcellular compartments, as derived from quantitative analysis of microalgal 3D models. Three cells are considered for every taxon or species. **(A)** Hexagons: *Micromonas*; circles: *Pelagomonas*; stars: *Emiliania*; squares: *Galdieria*; triangles: *Phaeodactylum*; diamonds: *Symbiodinium.* **(B)** Because of the much larger size of *Symbiodinium* cells, all the other taxa are compacted in a bottom left cluster in the plot Linear regressions between *Symbiodinium* cells and this cluster can be easily found, biasing the overall correlation analysis.

**Supplementary Figure 3.**
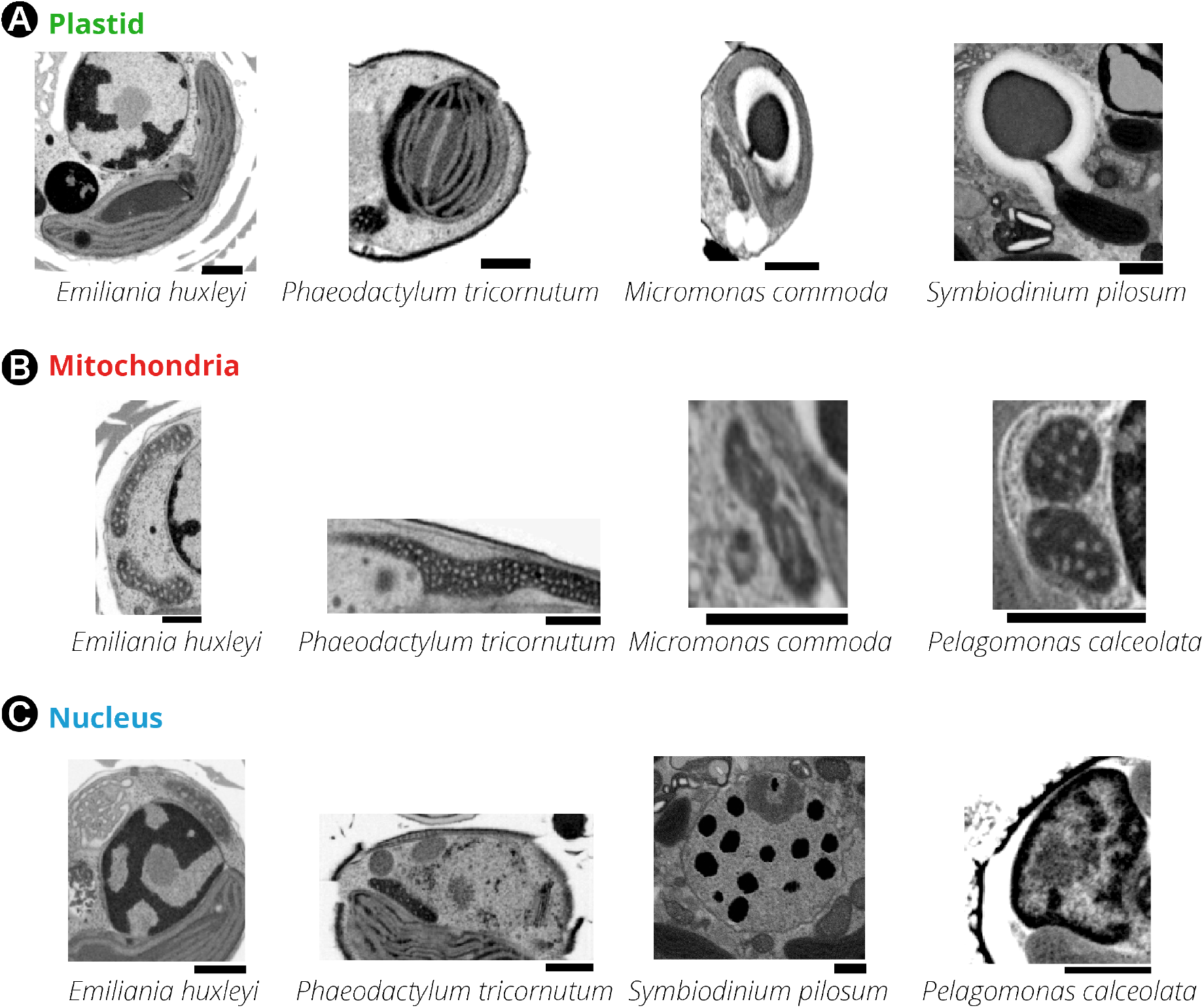
Subcellular features of different phytoplankton taxa: 2D FIB-SEM frames of pyrenoids (**A**), mitochondria (**B**) and nuclei (**C**). Scale bar: 500 nm.

**Supplementary Figure 4.**
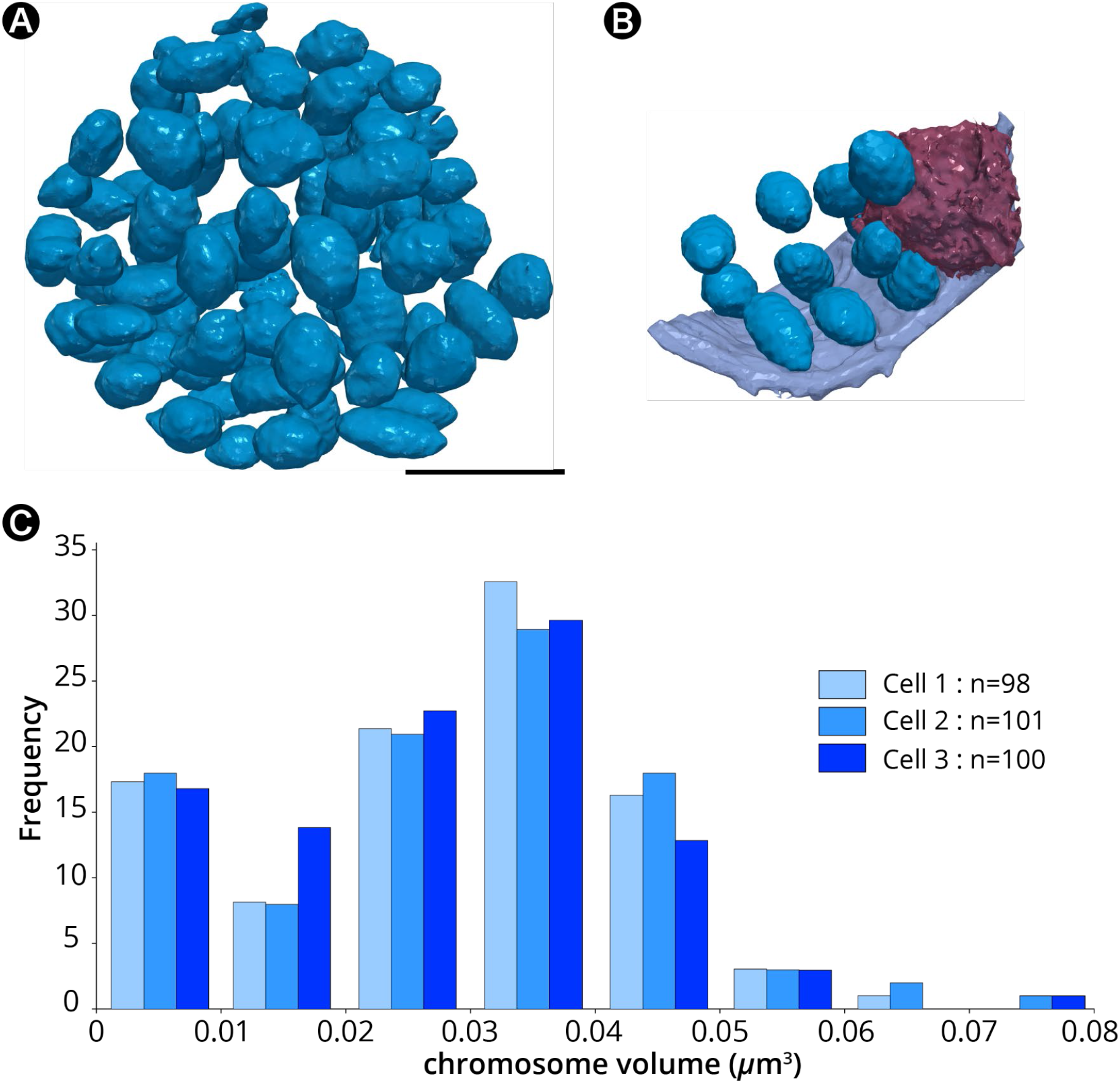
Topological arrangement of chromosomes inside the nucleus of the dinoflagellate Symbiodinium pilosum. **A:** 3D arrangement of chromosomes (blue) inside the nucleus. **B:** Contact points with the nuclear envelope (purple) were detected for peripheral chromosomes (Bhaud et al., 2000) and for the nucleolus (wine). **C:** Chromosomes number and volume distribution in three cells of *Symbiodinium pilosum*. Scale bar: 1 μm.

***Supplementary table 4 quantitative analysis of the morphological features of different phytoplankton cells***

***Supplementary Video 1. Focus Ion Beam Scanning Electron Microscopy (FIB-SEM) based 3D reconstruction of a whole cell of* Micromonas commoda.**

***Supplementary Video 2. Focus Ion Beam Scanning Electron Microscopy (FIB-SEM) based 3D reconstruction of a whole cell of* Pelagomonas calceolata.**

***Supplementary Video 3. Focus Ion Beam Scanning Electron Microscopy (FIB-SEM) based 3D reconstruction of a whole cell of* Emiliania huxleyi.**

***Supplementary Video 4. Focus Ion Beam Scanning Electron Microscopy (FIB-SEM) based 3D reconstruction of a whole cell of* Galdieria sulphuraria.**

***Supplementary Video 5. Focus Ion Beam Scanning Electron Microscopy (FIB-SEM) based 3D reconstruction of a whole cell of* Phaeodactylum tricornutum.**

***Supplementary Video 6. Focus Ion Beam Scanning Electron Microscopy (FIB-SEM) based 3D reconstruction of a whole cell of* Symbiodinium pilosum.**

***Supplementary Video 7. 3D representation of Focus Ion Beam Scanning Electron Microscopy (FIB-SEM) based 3D reconstruction of a whole cell of* Emiliania huxleyi**.

Grey: coccosphere (exoskeleton); blue: nucleus (dark blue: heterochromatin; light blue: euchromatin; violet: nucleolus); green: plastid; red: mitochondria; purple: Golgi apparatus and associated coccolith vesicles; yellow: storage compartments.

**Supplementary Table 1.** Information on the phytoplankton taxa analysed in this study (class, species and cell size or diameter).

| Class | Species | Cell size |
| --- | --- | --- |
| Bacillariophyceae | *Phaeodactylum tricornutum* Pt1 8.6 | 13-15 µm of length |
| Pelagophyceae | *Pelagomonas calceolata* RCC 100 | ≤ 3 µm in diameter |
| Dinophyceae | *Symbiodinium pilosum* (clade A) RCC 4014 | 7-8 µm in diameter |
| Prymnesiophyceae | *Emiliana huxleyi* RCC 909 | 3-8 µm in diameter |
| Mammiellophyceae | *Micromonas commoda* RCC 827 | < 2 µm in diameter |
| Cyanidiophyceae | *Galdieria sulphuraria* SAG21.92 | 3-9 µm |

**Supplementary Table 2.** Comparison of standard 3D mesh formats in MeshLab in the case of Symbiodinium pilosum cells. MeshLab is able to read almost all mesh formats (.obj;.stl;.ply) but not the.vtk format generated by 3Dslicer. While both the .obj and .ply files are smaller in size (and therefore easier to handle for visualization and animation), MeshLab encounter watertight problems with these formats (**: mesh surface is not closed).Therefore, quantitative analyses of volumes and surfaces were not possible unless using the .stl file format. After remeshing, Meshlab successfully reduced the number of polygons contained in the various objects, thereby generating smaller files that were easier to handle because of their reduced memory footprint. In addition, we checked that this mesh simplification procedure did not alter the volumetric information.

| 3D output formats from 3D Slicer | .stl format* |  | .obj format* |  | .ply format* |  | .vtk format* |  |
| --- | --- | --- | --- | --- | --- | --- | --- | --- |
|  | Before remesh | After remesh | Before remesh | After remesh | Before remesh | After remesh | Before remesh | After remesh |
| 3D model file size | 583.4 Mb | 100 Mb | 1 Gb | 178.9 Mb | 223.6 Mb | 36.6 Mb | 256.6 Mb | - |
| Model polygons | 7816626 | 2000000 | 7816626 | 2000000 | 7816626 | 2000000 | - | - |
| Volume information | 201.5 $\mu\text{m}^3$ | 201.5 $\mu\text{m}^3$ | Watertight problem** | - | Watertight problem** | - | - | - |

**Supplementary Table 3:** *The volume and surface of 3D models of plastids in* Emiliania huxleyi. The volume and surface were computed using 3D Slicer software and compared with those obtained using the STL python package and MeshLab software.

|  |  |  |  |
| --- | --- | --- | --- |
| Geometrical analysis | 3D Slicer (nrrd file) | Python based computing (STL file) | MeshLab (STL file) |
| Vol. model test ($\mu\text{m}^3$) | 1.166 | 1.173 | 1.182 |
| Surf. model test ($\mu\text{m}^2$) | Not provided | 20.992 | 20.89 |

## References

Ahrens J., B. Geveci, and C. Law. 2005. ParaView: an end-user tool for large data visualization. The visualization handbook. 717.

Allen, M.B. 1959. Studies with Cyanidium caldarium, an anomalously pigmented chlorophyte. Archiv fur Mikrobiologie. 32:270–277.

Andersen, R.A., J.C. Bailey, J. Decelle, and I. Probert. 2015. Phaeocystis rex sp. nov. (Phaeocystales, Prymnesiophyceae): a new solitary species that produces a multilayered scale cell covering. European Journal of Phycology. 50:207–222.

Andersen, R.A., G.W. Saunders, M.P. Paskind, and J.P. Sexton. 1993. Ultrastructure and 18s Ribosomal-Rna Gene Sequence for Pelagomonas-Calceolata Gen Et Sp-Nov and the Description of a New Algal Class, the Pelagophyceae Classis Nov. J Phycol. 29:701–715.

Badger, M.R., and G.D. Price. 1994. The Role of Carbonic Anhydrase in Photosynthesis. Annual Review of Plant Physiology and Plant Molecular Biology. 45:369–392.

Bailleul B., N. Berne, O. Murik, D. Petroutsos, J. Prihoda, A. Tanaka, V. Villanova, R. Bligny, S. Flori, D. Falconet, A. Krieger-Liszkay, S. Santabarbara, F. Rappaport, P. Joliot, L. Tirichine, P.G. Falkowski, P. Cardol, C. Bowler, and G. Finazzi. 2015. Energetic coupling between plastids and mitochondria drives CO_2_ assimilation in diatoms. Nature. 524:366–369.

Berges J., D. Franklin, and P. Harrison. 2004. Evolution of an artificial seawater medium: Improvements in enriched seawater, artificial water over the last two decades (Vol. 37: 1138–1145). J Phycol. 40:619.

Beuvier T., I. Probert, L. Beaufort, B. Suchéras-Marx, Y. Chushkin, F. Zontone, and A. Gibaud. 2019. X-ray nanotomography of coccolithophores reveals that coccolith mass and segment number correlate with grid size. Nature communications. 10:1–8.

Bhaud Y., D. Guillebault, J. Lennon, H. Defacque, M.O. Soyer-Gobillard, and H. Moreau. 2000. Morphology and behaviour of dinoflagellate chromosomes during the cell cycle and mitosis. Journal of cell science. 113 (Pt 7):1231–1239.

Blank, R.J. 1987. Cell Architecture of the Dinoflagellate Symbiodinium Sp Inhabiting the Hawaiian Stony Coral Coral Montipora-Verrucosa. Mar Biol. 94:143–155.

Cignoni P., M. Callieri, M. Corsini, M. Dellepiane, F. Ganovelli, and G. Ranzuglia. 2008. MeshLab: an Open-Source Mesh Processing Tool. Sixth Eurographics Italian Chapter Conference:129–136.

Colin S., L.P. Coelho, S. Sunagawa, C. Bowler, E. Karsenti, P. Bork, R. Pepperkok, and C. de Vargas. 2017. Quantitative 3D-imaging for cell biology and ecology of environmental microbial eukaryotes. eLife. 6.

Dauvillee D., P. Deschamps, J.P. Ral, C. Plancke, J.L. Putaux, J. Devassine, A. Durand-Terrasson, A. Devin, and S.G. Ball. 2009. Genetic dissection of floridean starch synthesis in the cytosol of the model dinoflagellate Crypthecodinium cohnii. Proceedings of the National Academy of Sciences of the United States of America. 106:21126–21130.

de Vargas C., S. Audic, N. Henry, J. Decelle, F. Mahe, R. Logares, E. Lara, C. Berney, N. Le Bescot, I. Probert, M. Carmichael, J. Poulain, S. Romac, S. Colin, J.M. Aury, L. Bittner, S. Chaffron, M. Dunthorn, S. Engelen, O. Flegontova, L. Guidi, A. Horak, O. Jaillon, G. Lima-Mendez, J. Lukes, S. Malviya, R. Morard, M. Mulot, E. Scalco, R. Siano, F. Vincent, A. Zingone, C. Dimier, M. Picheral, S. Searson, S. Kandels-Lewis, C. Tara Oceans, S.G. Acinas, P. Bork, C. Bowler, G. Gorsky, N. Grimsley, P. Hingamp, D. Iudicone, F. Not, H. Ogata, S. Pesant, J. Raes, M.E. Sieracki, S. Speich, L. Stemmann, S. Sunagawa, J. Weissenbach, P. Wincker, and E. Karsenti. 2015. Ocean plankton. Eukaryotic plankton diversity in the sunlit ocean. Science. 348:1261605.

Decelle J., S. Romac, R.F. Stern, E.M. Bendif, A. Zingone, S. Audic, M.D. Guiry, L. Guillou, D. Tessier, and Le Gall. 2015. Phyto REF: a reference database of the plastidial 16S rRNA gene of photosynthetic eukaryotes with curated taxonomy. Molecular ecology resources. 15:1435–1445.

Decelle J., H. Stryhanyuk, B. Gallet, G. Veronesi, M. Schmidt, S. Balzano, S. Marro, C. Uwizeye, P.H. Jouneau, J. Lupette, J. Jouhet, E. Marechal, Y. Schwab, N.L. Schieber, R. Tucoulou, H. Richnow Finazzi, and N. Musat. 2019. Algal Remodeling in a Ubiquitous Planktonic Photosymbiosis. Current biology: CB. 29:968–978 e964.

Decelle J., G. Veronesi, B. Gallet, H. Stryhanyuk, P. Benettoni, M. Schmidt, R. Tucoulou, M. Passarelli, S. Bohic, and P. Clode. 2020. Subcellular Chemical Imaging: New Avenues in Cell Biology. Trends in Cell Biology. 30.

del Pino Plumed M., A. Villarejo, A. de los Róos, G. García-Reina, and Z. Ramazanov. 1996. The CO 2- concentrating mechanism in a starchless mutant of the green unicellular alga Chlorella pyrenoidosa. Planta. 200:28–31.

Embleton, K.V., C.E. Gibson, and S.I. Heaney. 2003. Automated counting of phytoplankton by pattern recognition: a comparison with a manual counting method. J Plankton Res. 25:669–681.

Engel, B.D., M. Schaffer, L. Kuhn Cuellar, E. Villa, J.M. Plitzko, and W. Baumeister. 2015. Native architecture of the Chlamydomonas chloroplast revealed by in situ cryo-electron tomography. eLife. 4.

Field, C.B., M.J. Behrenfeld, J.T. Randerson, and P. Falkowski. 1998. Primary production of the biosphere: integrating terrestrial and oceanic components. Science. 281:237–240.

Flori S., P.H. Jouneau, B. Bailleul, B. Gallet, L.F. Estrozi, C. Moriscot, O. Bastien, S. Eicke, A. Schober, C.R. Bartulos, E. Marechal, P.G. Kroth, D. Petroutsos, S. Zeeman, C. Breyton, G. Schoehn, D. Falconet, and G. Finazzi. 2017. Plastid thylakoid architecture optimizes photosynthesis in diatoms. Nature communications. 8:15885.

Flori S., P.H. Jouneau, G. Finazzi, E. Marechal, and D. Falconet. 2016. Ultrastructure of the Periplastidial Compartment of the Diatom Phaeodactylum tricornutum. Protist. 167:254–267.

Gal A., A. Sorrentino, K. Kahil, E. Pereiro, D. Faivre, and A. Scheffel. 2018. Native-state imaging of calcifying and noncalcifying microalgae reveals similarities in their calcium storage organelles. Proceedings of the National Academy of Sciences of the United States of America. 115:11000–11005.

Gavelis, G.S., M. Herranz, K.C. Wakeman, C. Ripken, S. Mitarai, G.H. Gile, P.J. Keeling, and B.S. Leander. 2019. Dinoflagellate nucleus contains an extensive endomembrane network, the nuclear net. Scientific reports. 9:839.

Hawes C., and E. Hummel. 2015. Three-dimensional SEM - the future of cell imaging. Journal of microscopy. 259:79.

Helle, S.C., G. Kanfer, K. Kolar, A. Lang, A.H. Michel, and B. Kornmann. 2013. Organization and function of membrane contact sites. Biochimica et biophysica acta. 1833:2526–2541.

Hense, B.A., P. Gais, U. Jutting, H. Scherb, and K. Rodenacker. 2008. Use of fluorescence information for automated phytoplankton investigation by image analysis. J Plankton Res. 30:587–606.

Intergovernmental Panel on Climate, C. 2014. Climate Change 2014 – Impacts, Adaptation and Vulnerability: Part B: Regional Aspects: Working Group II Contribution to the IPCC Fifth Assessment Report: Volume 2: Regional Aspects. Cambridge University Press, Cambridge.

Itakura, A.K., K.X. Chan, N. Atkinson, L. Pallesen, L. Wang, G. Reeves, W. Patena, O. Caspari, R. Roth, and U. Goodenough. 2019. A Rubisco-binding protein is required for normal pyrenoid number and starch sheath morphology in Chlamydomonas reinhardtii. Proceedings of the National Academy of Sciences. 116:18445–18454.

Kikinis R., S.D. Pieper, and K.G. Vosburgh. 2014. 3D Slicer: a platform for subject-specific image analysis, visualization, and clinical support. In Intraoperative imaging and image-guided therapy. Springer. 277–289.

Kim J., M. Fabris, G. Baart, M.K. Kim, A. Goossens, W. Vyverman, P.G. Falkowski, and D.S. Lun. 2016. Flux balance analysis of primary metabolism in the diatom Phaeodactylum tricornutum. The Plant journal: for cell and molecular biology. 85:161–176.

Lopes Dos Santos A., T. Pollina, P. Gourvil, E. Corre, D. Marie, J.L. Garrido, F. Rodriguez, M.H. Noel, D. Vaulot, and W. Eikrem. 2017. Chloropicophyceae, a new class of picophytoplanktonic prasinophytes. Scientific reports. 7:14019.

Lupette J., A. Jaussaud, K. Seddiki, C. Morabito, S. Brugiere, H. Schaller, M. Kuntz, J.L. Putaux, P.H. Jouneau, F. Rebellle, D. Falconet, Y. Coute, J. Jouhet, M. Tardif, J. Salvaing, and E. Marechal. 2019. The architecture of lipid droplets in the diatom Phaeodactylum tricornutum. Algal Res. 38.

Meyer, M.T., A.J. McCormick, and H. Griffiths. 2016. Will an algal CO_2_-concentrating mechanism work in higher plants? Current opinion in plant biology. 31:181–188.

Meyer, M.T., C. Whittaker, and H. Griffiths. 2017. The algal pyrenoid: key unanswered questions. Journal of experimental botany. 68:3739–3749.

Moroney, J.V., and C.B. Mason. 1991. The Role of the Chloroplast in Inorganic Carbon Acquisition by Chlamydomonas-Reinhardtii. Can J Bot. 69:1017–1024.

Mueller-Schuessele, S.J., and M. Michaud. 2018. Plastid Transient and Stable Interactions with Other Cell Compartments. Methods in molecular biology. 1829:87–109.

Mustardy L., and G. Garab. 2003. Granum revisited. A three-dimensional model--where things fall into place. Trends in plant science. 8:117–122.

Narayan K., and S. Subramaniam. 2015. Focused ion beams in biology. Nature methods. 12:1021–1031.

Not F., R. Siano, W.H.C.F. Kooistra, N. Simon, D. Vaulot, and I. Probert. 2012. Diversity and ecology of eukaryotic marine phytoplankton. In Advances in botanical research 64:1–53.

Oliphant, T.E. 2007. Python for scientific computing. Computing in Science & Engineering. 9:10–20.

Ramazanov Z., M. Rawat, M.C. Henk, C.B. Mason, S.W. Matthews, and J.V. Moroney. 1994. The induction of the CO_2_-concentrating mechanism is correlated with the formation of the starch sheath around the pyrenoid of Chlamydomonas reinhardtii. Planta. 195:210–216.

Ranzuglia G., M. Callieri, M. Dellepiane, P. Cignoni, and R. Scopigno. 2012. MeshLab as a complete tool for the integration of photos and color with high resolution 3D geometry data. CAA 2012 Conference Proceedings:406–416.

Rodenacker K., B. Hense, U. Jutting, and P. Gais. 2006. Automatic analysis of aqueous specimens for phytoplankton structure recognition and population estimation. Microsc Res Techniq. 69:708–720.

Russo, F. 2002. An image enhancement technique combining sharpening and noise reduction. IEEE Transactions on Instrumentation and Measurement. 51:824–828.

Sartori A., R. Gatz, F. Beck, A. Rigort, W. Baumeister, and J.M. Plitzko. 2007. Correlative microscopy: bridging the gap between fluorescence light microscopy and cryo-electron tomography. Journal of structural biology. 160:135–145.

Schulze K., U.M. Tillich, T. Dandekar, and M. Frohme. 2013. PlanktoVision - an automated analysis system for the identification of phytoplankton. Bmc Bioinformatics. 14.

Scorrano L., M.A. De Matteis, S. Emr, F. Giordano, G. Hajnoczky, B. Kornmann, L.L. Lackner, T.P. Levine, L. Pellegrini, K. Reinisch, R. Rizzuto, T. Simmen, H. Stenmark, C. Ungermann, and M. Schuldiner. 2019. Coming together to define membrane contact sites. Nature communications. 10:1287.

Sosik, H.M., and R.J. Olson. 2007. Automated taxonomic classification of phytoplankton sampled with imaging-in-flow cytometry. Limnology and Oceanography: Methods. 5:204–216.

Stephens, D.J., and V.J. Allan. 2003. Light microscopy techniques for live cell imaging. Science. 300:82–86.

Sviben S., A. Gal, M.A. Hood, L. Bertinetti, Y. Politi, M. Bennet, P. Krishnamoorthy, A. Schertel, R. Wirth, A. Sorrentino, E. Pereiro, D. Faivre, and A. Scheffel. 2016. A vacuole-like compartment concentrates a disordered calcium phase in a key coccolithophorid alga. Nature communications. 7:11228.

Taylor, A.R., C. Brownlee, and G. Wheeler. 2017. Coccolithophore cell biology: chalking up progress. Annual review of marine science. 9:283–310.

Tilokani L., S. Nagashima, V. Paupe, and J. Prudent. 2018. Mitochondrial dynamics: overview of molecular mechanisms. Essays in biochemistry. 62:341–360.

Titze B., and C. Genoud. 2016. Volume scanning electron microscopy for imaging biological ultrastructure. Biology of the cell. 108:307–323.

Toyokawa C., T. Yamano, and H. Fukuzawa. 2020. Pyrenoid Starch Sheath Is Required for LCIB Localization and the CO_2_-Concentrating Mechanism in Green Algae. Plant physiology. 182:1883.

Van Thinh L., D.J. Griffiths, and H. Winsor. 1986. Ultrastructure of Symbiodinium microadriaticum (Dinophyceae) symbiotic with Zoanthus sp. (Zoanthidea). Phycologia. 25:178–184.

Vaulot D., F.L. Gall, D. Marie, L. Guillou, and F. Partensky. 2004. The Roscoff Culture Collection (RCC): a collection dedicated to marine picoplankton. Nova Hedwigia. 79:49–70.

Villarejo A., F. Martinez, M.D. Plumed, and Z. Ramazanov. 1996. The induction of the CO_2_ concentrating mechanism in a starch-less mutant of Chlamydomonas reinhardtii. Physiol Plantarum. 98:798–802.

Viola R., P. Nyvall, and M. Pedersen. 2001. The unique features of starch metabolism in red algae. Proceedings of the Royal Society of London. Series B: Biological Sciences. 268:1417–1422.

Wietrzynski W., M. Schaffer, D. Tegunov, S. Albert, A. Kanazawa, J.M. Plitzko, W. Baumeister, and B.D. Engel. 2020. Charting the native architecture of Chlamydomonas thylakoid membranes with single-molecule precision. eLife. 9:e53740.

Winder M., and U. Sommer. 2012. Phytoplankton response to a changing climate. Hydrobiologia. 698:5–16.

Zhang C., and T. Chen. 2001. Efficient feature extraction for 2D/3D objects in mesh representation. In Proceedings 2001 International Conference on Image Processing (Cat. No. 01CH37205). Vol. 3. IEEE. 935–938.

